# High-throughput ANI Analysis of 90K Prokaryotic Genomes Reveals Clear Species Boundaries

**DOI:** 10.1101/225342

**Authors:** Chirag Jain, Luis M. Rodriguez-R, Adam M. Phillippy, Konstantinos T. Konstantinidis, Srinivas Aluru

**Affiliations:** School of Computational Science and Engineering, Georgia Institute of Technology, Atlanta GA 30332; School of Civil and Environmental Engineering, Georgia Institute of Technology, Atlanta GA 30332; School of Biological Sciences, Georgia Institute of Technology, Atlanta GA 30332; National Human Genome Research Institute, National Institutes of Health, Bethesda MD 20894; Institute for Data Engineering and Science, Georgia Institute of Technology, Atlanta GA 30332

**Keywords:** species concept, nucleotide identity, minhash, prokaryotic diversity

## Abstract

A fundamental question in microbiology is whether there is a continuum of genetic diversity among genomes or clear species boundaries prevail instead. Answering this question requires robust measurement of whole-genome relatedness among thousands of genomes and from diverge phylogenetic lineages. Whole-genome similarity metrics such as Average Nucleotide Identity (ANI) can provide the resolution needed for this task, overcoming several limitations of traditional techniques used for the same purposes. Although the number of genomes currently available may be adequate, the associated bioinformatics tools for analysis are lagging behind these developments and cannot scale to large datasets. Here, we present a new method, FastANI, to compute ANI using alignment-free approximate sequence mapping. Our analyses demonstrate that FastANI produces an accurate ANI estimate and is up to three orders of magnitude faster when compared to an alignment (e.g., BLAST)-based approach. We leverage FastANI to compute pairwise ANI values among all prokaryotic genomes available in the NCBI database. Our results reveal a clear genetic discontinuity among the database genomes, with 99.8% of the total 8 billion genome pairs analyzed showing either >95% intra-species ANI or <83% inter-species ANI values. We further show that this discontinuity is recovered with or without the most frequently represented species in the database and is robust to historic additions in the public genome databases. Therefore, 95% ANI represents an accurate threshold for demarcating almost all currently named prokaryotic species, and wide species boundaries may exist for prokaryotes.

---

Large collections of prokaryotic genomes with varied ecologic and evolutionary histories are now publicly available. This deluge of genomic data provides the opportunity to more robustly evaluate important questions in microbial ecology and evolution, as well as underscores the need to advance existing bioinformatics approaches for the analysis of such big genomic data. One such question is whether bacteria (and other microbes) form discrete clusters (species), or, due to high frequency of horizontal gene transfer (HGT) and slow decay kinetics, a continuum of genetic diversity is observed instead. Studies based on a small number of closely related genomes have shown that genetic continuum may prevail [e.g., (1)]. On the other hand, other studies have argued that HGT may not be frequent enough to distort species boundaries, or that organisms within species exchange DNA more frequently compared to organisms across species, thus maintaining distinct clusters [e.g., (2)]. An important criticism of all these studies is that they have typically been performed with isolated genomes in the laboratory that may not adequately represent natural diversity due to cultivation biases, or were based on a small number of available genomes from a few phylogenetic lineages, which does not allow for robust conclusions to emerge. Therefore, it is still unclear if well-defined clusters of genomes are evident among prokaryotes and how to recognize them. Defining species is not only an important academic exercise but also has major practical consequences. For instance, the diagnosis of disease agents, the regulation of which organisms can be transported across countries and which organisms should be under quarantine, or the communication about which organisms or mixtures of organisms are beneficial to human, animals or plants, are all deeply-rooted on how species are defined.

One fundamental task in assessing species boundaries is the estimation of the genetic relatedness between two genomes. In recent years, the whole-genome average nucleotide identity (ANI) has emerged as a robust method for this task, with organisms belonging to the same species typically showing ≥95% ANI among themselves (3, 4). ANI represents the average nucleotide identity of all orthologous genes shared between any two genomes and offers robust resolution between strains of the same or closely related species (i.e., showing 80-100% ANI). The ANI measure does not strictly represent core genome evolutionary relatedness, as orthologous genes can vary widely between pairs of genomes compared. Nevertheless, it closely reflects the traditional microbiological concept of DNA-DNA hybridization relatedness for defining species (3), as it takes into account the fluid nature of the bacterial gene pool and hence implicitly considers shared function. Compared to sequencing of 16S rRNA genes, another highly popular, alternative traditional method for defining species and assessing their evolutionary uniqueness, ANI offers several important advantages such as higher resolution among closely related genomes. Finally, ANI can be estimated among draft (incomplete) genome sequences recovered from the environment using metagenomic or singe-cell techniques that do not encode a universally conserved gene such as the 16S rRNA gene (e.g., due to mis-assembly) but encode at least a few hundred shared genes, greatly expanding the number of sequences that can be studied and classified compared to a universal gene-based approach. Accordingly, ANI has been recognized internationally for its potential for replacing DNA-DNA hybridization as the standard measure of relatedness, as it is easier to estimate and represents portable and reproducible data (5, 6). Despite its strengths, traditional ANI calculation is based on alignment-based searches [e.g., BLAST (7)] and thus, remains computationally expensive due to the quadratic time complexity of alignment algorithms, and does not scale well with an increasing number of genomes.

### Significance Statement

With the advent of big data in genomics, fundamental biological questions can now be answered through data-driven approaches and scalable bioinformatics algorithms. Doing so, requires new algorithms that employ intelligent tradeoffs between computational speed and accuracy. This work presents such a scalable algorithm for computing the whole-genome genetic relatedness among genomes. By applying this algorithm to a large collection of prokaryotic genomes, our study addressed the following biological question: do well-defined clusters of genomes (species) exist?

Several variations of the original ANI calculation algorithm have been proposed (8–10), but these mainly modify the specific approach to identify shared genes and do not speed up the calculation substantially since they are all alignment-based. Accordingly, it is nearly impossible to calculate ANI values among the available microbial genomes to date, in the order of a hundred thousand, based on these approaches and commonly available computational resources. Importantly, the available genomic data is estimated to be a small fraction of the extant prokaryotic diversity (11), and the number of new genomes determined continues to grow exponentially. Therefore, new computational solutions are needed to scale-up with the available and forthcoming data.

A couple of such solutions have been proposed recently, borrowing concepts from ‘big data’ analysis in other scientific domains. MinHash is a technique for quick estimation of similarity of two sets, initially developed for the detection of near-duplicate web documents in search engines at the scale of the World Wide Web (12). Recently, this technique was successfully adapted for designing new fast algorithms in bioinformatics such as for genome assembly (13, 14) and long read mapping problems (15). Ondov *et. al*. (16) provided the first proof-of-concept implementation called Mash for fast estimation of ANI using this technique. Even though Mash has been reported to be multiple orders of magnitude faster than alignment-based ANI computation, a straight-forward adoption of the MinHash technique to the problem of computing ANI has been found to be inaccurate for incomplete draft genomes (17). Further, there is a limit on how well Mash can approximate ANI especially for moderately divergent genomes (e.g., showing 80-90% ANI), as Mash similarity measurement is not restricted to the shared genomic regions, whereas ANI considers only the shared genome.

In this study, we alleviate the computational bottleneck in ANI computation by developing FastANI, a novel algorithm utilizing Mashmap (15) as its MinHash based alignment-free sequence mapping engine. FastANI provides ANI values that are essentially identical to the alignment-based ANI values for both complete and draft quality genomes that are related in the 80% to 100% nucleotide identity range. Therefore, FastANI should enable the accurate estimation of pairwise ANI values for large cohorts of genomes or evaluation of the novelty of a query draft genome by comparing it against the full collection of available prokaryotic genomes.

## Results

We developed FastANI, an algorithm for efficiently computing pairwise average nucleotide identities among a large set of genomes, and applied it to determine whether genomic data supports the existence of genetic discontinuity and clear special boundaries among prokaryotic species. We first demonstrate FastANI yields accuracy on par with the widely accepted BLASTn based approaches, and then leverage its computational efficiency to analyze genomic relatedness within and across species.

### Datasets

To test accuracy and speed, we evaluated FastANI on both high-quality closed genomes from NCBI RefSeq database as well as draft genome assemblies downloaded from the prokaryote section of the NCBI Genome database. We first removed poor quality genome assemblies with low N50 length (< 10 Kbp). In total, four datasets were used (D1 through D4). Dataset D1 is the set of closed prokaryotic genomes downloaded from RefSeq database. Datasets D2, D3, and D4 are collections of draft genome assemblies of *Bacillus cereus, Escherichia coli* and *Bacillus anthracis*, respectively. These sizable datasets represent genomes showing different levels of identity among themselves and varying values of completeness and assembly quality. For each dataset, one genome was selected as the query genome and its ANI was computed with every genome in the complete dataset (see Table 1).

**Table 1.** Datasets used for testing accuracy and speed of FastANI.

| Id | Reference clade | No. of Genomes | Median N50 (Mbp) | Query Genome |
| --- | --- | --- | --- | --- |
| D1 | NCBI RefSeq | 1,675 | 3.14 | *E. coli* K-12 MG1655 |
| D2 | *Bacillus cereus* | 570 | 1.16 | *B. anthracis* 52-G |
| D3 | *Escherichia coli* | 4,271 | 0.15 | *E. coli* strain |
| D4 | *Bacillus anthracis* | 464 | 0.59 | *B. anthracis* strain |

### Accuracy

We evaluated FastANI against the BLASTn based method of computing ANI (9), henceforth referred as ANI_b_, and the ANI values predicted by the Mash tool (16). User documentation for Mash recommends using larger sketch size (i.e., k-mer sample) than the default to obtain higher accuracy (16). Accordingly, we ran Mash with both the default sketch size of 1*K* as well as increase it up to 100*K*.

FastANI achieves near perfect linear correlation with ANI_b_ on all datasets D1-D4 (Figure 1). Mash results improve with increasing sketch size, particularly for D1. However, even when executed with the largest sketch size of 100K, Mash results diverge from ANI_b_ values on datasets D1, D3 and D4. For D1, this primarily appears to be caused by divergent genomes (e.g., showing < 90% ANI). For D3, Mash diverges on closely related genomes due to fragmented and incomplete genome assemblies of the draft genomes. Dataset D4 is challenging because its constituent genomes are closely related strains of *Bacillus anthracis*, with ANI_b_ > 99.9 for all the pairs. FastANI provides much better precision than Mash in D4 dataset, and therefore, can be used to discriminate between very closely related microbial strains such as those of different epidemic outbreaks. However, for two genomes out of the 464, FastANI estimates are diverging from ANI_b_. To investigate further, we visualized gene synteny pattern using Mauve (18) and found that these two genome sequences have many re-arrangements with respect to the query genome (Supplementary Fig. S1). Given that *B. anthracis* strains typically show high genome synteny (19), these results indicate that the two genomes were incorrectly assembled. Incorrect data will yield unpredictable results not only with FastANI but using any method that assesses genetic relatedness, including phylogeny-based methods. If the two incorrect *B. anthracis* assemblies are removed, FastANI’s correlation with ANI_b_ improves to 0.944 in D4.

**Fig. 1.**
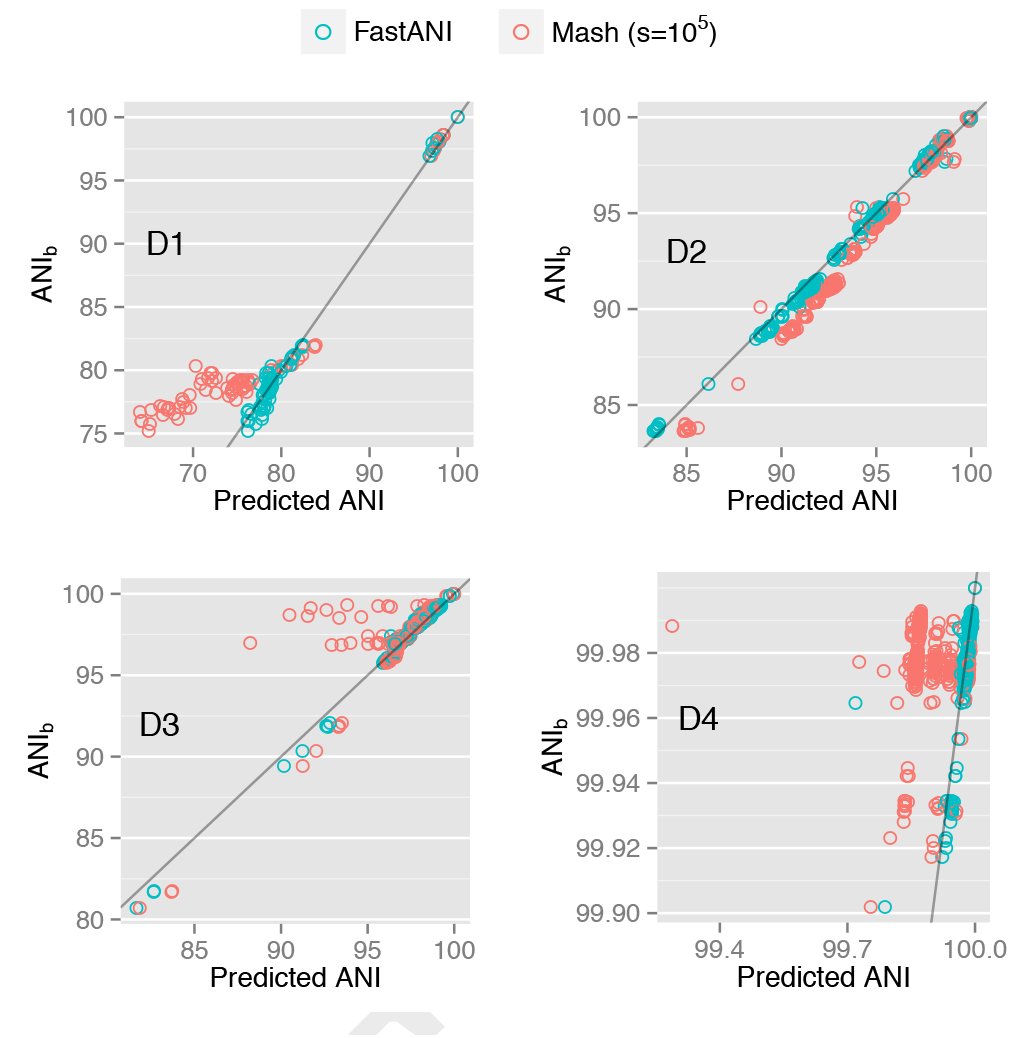
Plots showing how FastANI and Mash-based ANI (sketch size = 10^5^) output correlate with ANI_b_ values for datasets D1-D4. Because FastANI assumes a probabilistic identity cutoff that is set to 80% by default, it reports 76, 570, 4,271 and 464 genome matches for the individual queries in datasets D1-D4 respectively. To enable a direct quality comparison against FastANI, Mash is executed for only those pairs that are reported by FastANI. Notice that each dataset encompass a different nucleotide identity range (x-axes). Gray line represents a straight line *y = x* plot for reference. Pearson correlation coefficients corresponding to these plots are listed separately in Table 2.

These correlation results demonstrate FastANI provides significant quality improvement over Mash (see Table 2), and can be a reasonable substitute for ANI_b_. Aggregate results over all the datasets D1-D4 (Figure 2) led us to conclude that FastANI can tolerate variable assembly quality and completeness. Most importantly, it correlates well with ANI_b_ in the desired identity range 80% – 100%.

**Table 2.** Comparison of FastANI and Mash-based ANI accuracy by measuring their Pearson correlation coefficients with ANI_b_ values. Mash is executed with sketch sizes (-s): 1,000 (default), 10,000 and 100,000. FastANI achieves > 0.99 correlation with ANI_b_ on D1-D3. Its correlation value on D4 improves from 0.681 to 0.944 if the two incorrect assemblies present in D4 are not taken into account.

| Dataset | FastANI | Mash<br>-s 10<sup>3</sup> | Mash<br>-s 10<sup>4</sup> | Mash<br>-s 10<sup>5</sup> |
| --- | --- | --- | --- | --- |
| D1 | **0.995** | 0.594 | 0.932 | 0.935 |
| D2 | **0.999** | 0.996 | 0.997 | 0.997 |
| D3 | **0.995** | 0.944 | 0.944 | 0.944 |
| D4 | **0.681** | -0.040 | 0.003 | 0.010 |

**Fig. 2.**
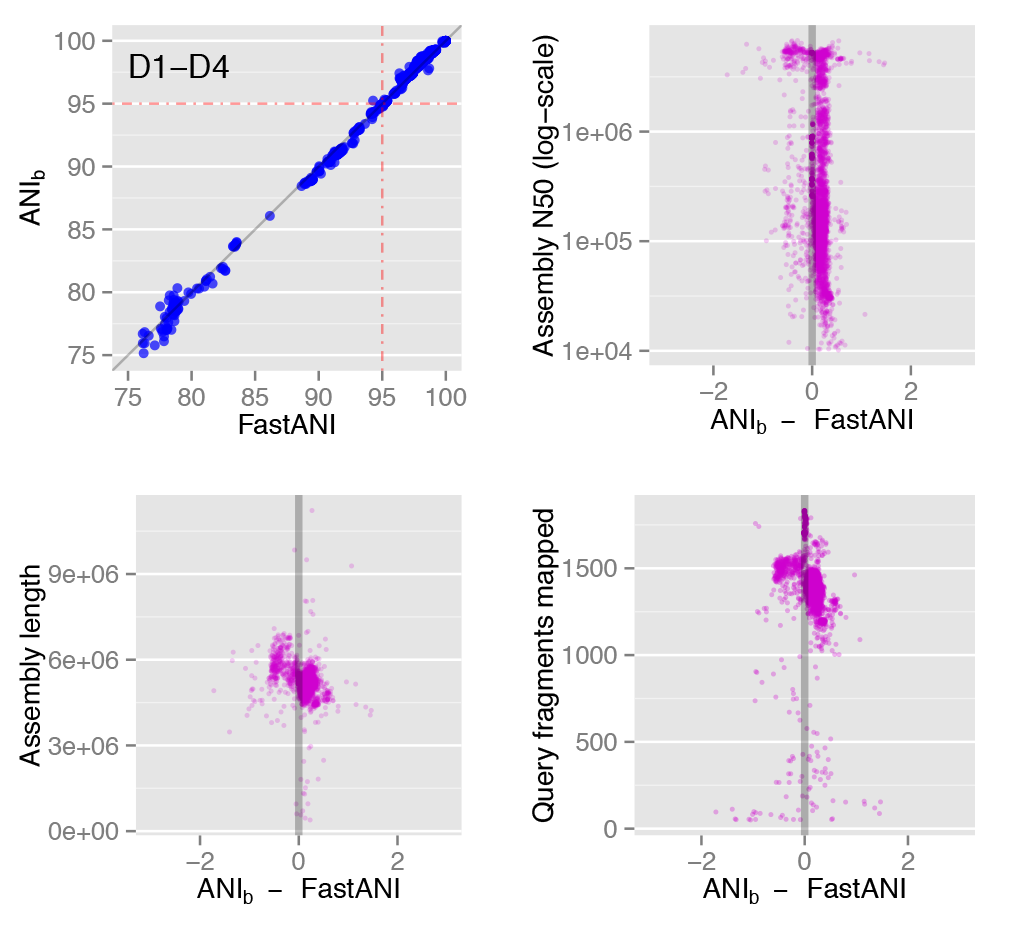
FastANI’s aggregate accuracy and error characteristics based on datasets D1-D4. Upper left plot shows the FastANI and ANI_b_ correlation. The remaining three plots show differences between FastANI and ANI_b_ value versus reference genome assembly quality (N50 and length) and the number of reciprocal fragments that matched between query and reference genome for each comparison. These results show no biases associated with these factors.

### Computational Speedup

FastANI is designed to efficiently process large assembly datasets with ordinary compute resources. For FastANI’s sequential and parallel runtime evaluation, we used a single compute node with two Intel Xeon E5-2698 v4 20-core processors. First we show runtime comparison of FastANI and ANI_b_ using serial execution (single thread, single process) using all datasets in Table 3. FastANI operation consists of indexing phase followed by compute phase, for which we measured the runtime separately. For any database, indexing all the reference genomes needs to be done only once, and thereafter, FastANI can compute ANI estimates for any number of input query genomes against the reference genomes. Therefore, speedup in Table 3 is measured with respect to FastANI compute time. We observe that the runtime improvement due to FastANI varied from 50x for D3 to 782x for D1. FastANI speed-up was the highest on D1 because NCBI Refseq database contains a diverse set of prokaryotic genomes. This is attributable to the fact that the algorithm underlying FastANI is able to prune distant genomes (ANI ≪ 80%) efficiently. On the contrary, ANI values for all genomes in datasets D2-D4 were high > 80%.

**Table 3.** Comparison of execution time of FastANI versus ANI_b_. Speedup in the last column is measured as the ratio of ANI_b_’s runtime and FastANI’s compute time.

| Dataset | FastANI |  | ANI<sub>b</sub> (sec) | Speedup |
| --- | --- | --- | --- | --- |
|  | Indexing (sec) | Compute (sec) |  |  |
| D1 | 468.2 | 16.76 | 13,113 | **782x** |
| D2 | 195.7 | 264.76 | 18,155 | **69x** |
| D3 | 1538 | 1980.8 | 99,317 | **50x** |
| D4 | 128.8 | 214.53 | 11,051 | **52x** |

To accelerate ANI computation even further, FastANI can be trivially parallelized using multi-core parallel execution. One way to achieve this is to split the reference genomes in several equal-size parts. This way, each instance of FastANI process can search query genome(s) against each part of the reference database independently. We utilized this scheme and evaluated scalability using up to 80 FastANI parallel processes. Compared to the sequential execution time listed in Table 3, runtime of the compute phase reduced to 2, 8, 46 and 6 seconds for datasets D1-D4 respectively (Figure 3). These results confirm FastANI can be used to query against databases containing thousands of genomes in a few seconds.

**Fig. 3.**
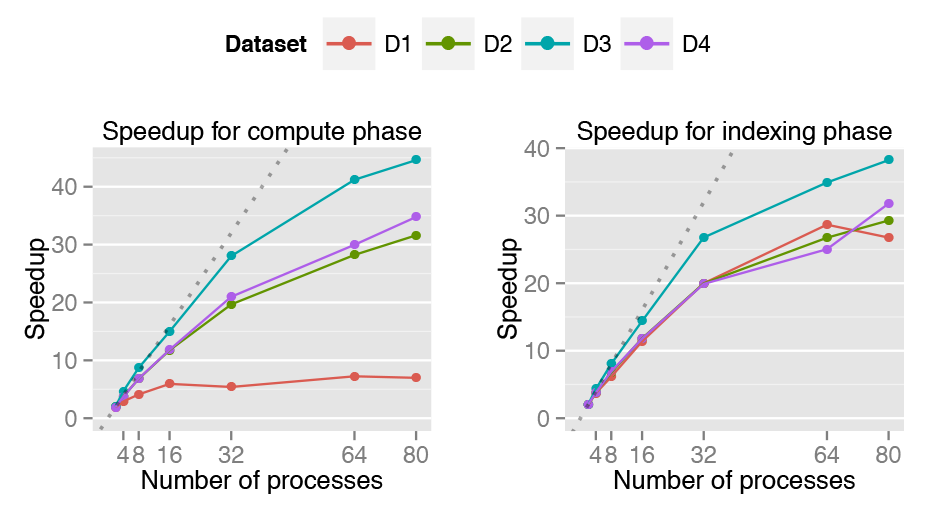
Scaling results of FastANI’s execution time using datasets D1-D4 on a compute node with 40 physical cores. We executed parallel FastANI processes where each process was assigned an equal sized random part of the reference database for computing ANI. Left and right plots evaluate FastANI’s compute and indexing phase, respectively. FastANI achieves reasonable speedups on all datasets except the compute phase in D1, as its runtime of 17s on a single core is too small to begin with (Table 3). FdastANI reduces this to 2 seconds using 80 parallel processes.

For the above experiments, FastANI required a maximum 48 GB memory for D3, our largest dataset for this experiment. For databases much larger than D3, peak memory usage can be reduced by either distributing the compute across multiple nodes in a cluster or processing chunks of the reference database one by one, as necessary.

### Large-scale Pairwise Comparison Indicates Genetic Discontinuity

We examined the distribution of pairwise ANI values between all 91,761 prokaryotic assemblies that existed in the NCBI Genome database as of March 15, 2017. Prior to analysis, we removed 2,262 genomes due to short N50 length (< 10 Kbp). In our analysis, the ANI between each pair of genomes A and B is computed twice, once with *A* as query genome and again with *B* as query genome. This choice did not meaningfully alter the ANI value reported by FastANI unless the draft genomes are incorrectly assembled or contaminated (Supplementary Fig. S2). Computing pairwise ANI values for the entire database took 77K CPU hours for all 8.01 billion comparisons. To our knowledge, this is the largest cohort of genomes for which ANI has been computed. The largest previously published ANI analysis included 86 million comparisons and took 190K CPU hours (20).

Among the total of 8.01 billion pairwise comparisons, 679,765,100 yielded ANI values in the 76-100% range. The distribution of these ANI values reveals a clear and wide discontinuity in the identity range of 83-95% (Figure 4a). FastANI reported only 17,132,536 ANI values (i.e., 2.5% of the 679,765,100 pairs) within the range of 83% to 95%. The genetic discontinuity was apparent even when all named species were randomly drawn with species-dependent probabilities that ensured the same expected representation of highly sampled and sparsely sampled species in the final set (data not shown).

**Fig. 4.**
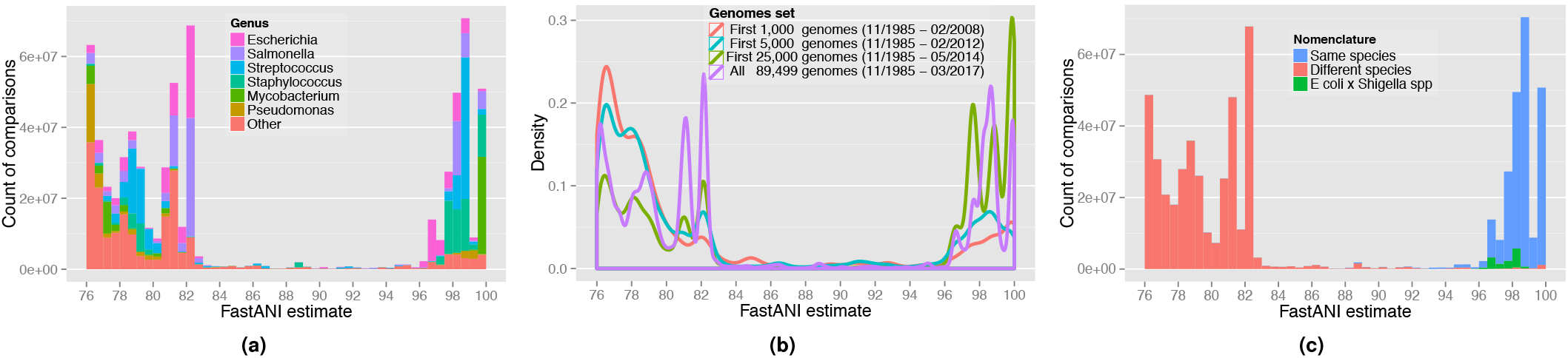
**a.** Histogram plot showing the distribution of ANI values among the 90K genomes. Only ANI values in the 76-100% range are shown. Out of total 8.01 billion pairwise genome comparisons, FastANI reported only 17M ANI values (0.21%) with ANI between 83% and 95% indicating a wide genetic discontinuum. Multiple colors are used to show how genomes from different genera are contributing to this distribution. Few peaks in the histogram arise from genera that have been extensively sequenced and dominate the database. **b.** Density curves of ANI values in the ANI range 76-100%. Each curve shows the density curve corresponding to the database at a particular time period. Wide discontinuity in all four curves is observed consistently, **c.** Distribution of ANI values with each comparison labeled by the nomenclature of genomes being compared. All the comparisons between *Escherichia coli* and *Shigella* spp. have been labeled separately. The 95% ANI threshold on x-axis serves as a valid classifier for comparisons belonging to same and different species.

The frequency of intra-vs. inter-species genomes sequenced in the NCBI database has changed over time, with earlier sequencing efforts targeting distantly related organisms in order to cover phylogenetic diversity while efforts in more recent years targeted more closely related organisms for microdiversity or epidemiological studies (Supplementary Fig. S3). We confirmed that discontinuity pattern has been maintained at different time points in the past (Figure 4b). In previous taxonomic studies, 95% ANI cutoff is the most frequently used standard for species demarcation. Density curves in the figure show that the two peaks consistently lie on either side of the 95% ANI value.

Finally, we tested the correlation between standing nomenclature and the 95% ANI demarcation. As per this standard, we should expect a pair of genomes to have ANI value ≥ 95% if and only if both genomes are members of same species. From the complete set of 89,499 genomes, we identified the subset for which we could determine the named species for each genome. Whenever available (9% of the total genomes), we recovered the links to NCBI taxonomy to determine the species. For the remainder of the genomes, we inferred the species from the organism name given in the GenBank file, excluding all entries with ambiguous terms (sp, cf, aff, bacterium, archeon, endosymbiont), resulting in the species-wise classification of an additional 78% of the genomes. The remainder 13% of the genomes were discarded.

We evaluated the distribution of ANI values in this subset in comparison to the named species that the corresponding genomes were assigned to (Figure 4c). The ≥ 95% ANI criterion reflects same named species with a recall frequency of 98.5% and a precision of 93.1%. We further explored the values affecting precision, i.e., 6.9% of ANI values above 95% that were obtained for genomes assigned to different named species. Among those, 5.6% are due to comparisons between *Escherichia coli* and *Shigella* spp., a case in which the inconsistency between taxonomy and genomic relatedness is well documented (1) (highlighted in green in Figure 4c). The remaining 1.3% of the cases mostly exist within the *Mycobacterium* genus (0.5%), which includes a group of closely related named species as part of the *M. tuberculosis* complex such as *M. tuberculosis* (reference), *M. canettii* (ANI 97-99% against reference), *M. bovis* (ANI 99.6%), *M. microti* (ANI 99.8-99.9%), and *M. africanum* (ANI 99.9%), among others. An additional 0.2% of the cases correspond to comparisons between *Neisseria gonorrhoeae* and *N. meningitidis*, two species with large representation in the database and ANI values close to 95% (Inter-quartile range: 94.9-95.2%). Excluding the cases of *E. coli* vs. *Shigella* alone, precision increases to 98.7%. With both recall and precision values ≥ 98.5%, these results corroborate the utility of ANI for species demarcation, which is consistent with previous studies based on a much smaller dataset of genomes (4, 20).

## Discussion

Our results indicate that FastANI robustly estimates ANI values between both complete and draft genomes while reducing the computing time by two to three orders of magnitude. We leveraged the computational efficiency offered by FastANI to evaluate the distribution of ANI values in a set of over 90,000 genomes, and demonstrate that genetic relatedness discontinuity can be consistently identified among these genomes around 95% ANI. This discontinuity is recovered with or without the most frequently represented species in the database, is robust to historic additions in the public databases, and it represents an accurate threshold for demarcating almost all currently named prokaryotic species.

Given also the large number of genomes used in our analysis that represented all major prokaryotic lineages, it is likely that the discontinuity represents a real biological signature and is not driven by cultivation or other biases. It is also important to note that these results are consistent with metagenomics analysis of natural microbial communities, which have showed that the communities are composed of predominantly sequence-discrete populations [reviewed in (21)]. The biological mechanisms underlying this genetic discontinuity are not clear but should be subject of future research for a more complete understanding of prokaryotic species. The mechanisms could involve a dramatic drop in recombination frequency around 90-95% ANI, which could account for the discontinuity if bacteria evolve sexually [reviewed in (22)], or ecological sweeps that remove diversity due to competition [reviewed in (23, 24)]. A genomic nucleotide diversity of 5-10% translates to tens of thousands of years of evolution time, which provides ample opportunities for ecological or genetic sweeps to occur. Nonetheless, the existence of genetic discontinuity represents a major finding that can help define species more accurately and has important practical consequences for recognizing and communicating about prokaryotic species.

As a general-purpose research tool, we expect FastANI to be useful for analysis of both clinical and environmental microbial genomes. It can be used for studying the inter- and intra-species diversity within large collections of genomes. It should also accelerate the study of the novelty of new species or phenotypic similarity of a query genome sequence in comparison to all available genomes.

## Materials and Methods

Before describing FastANI (Figure 5), we briefly review Mashmap that underlies the FastANI algorithm.

**Fig. 5.**
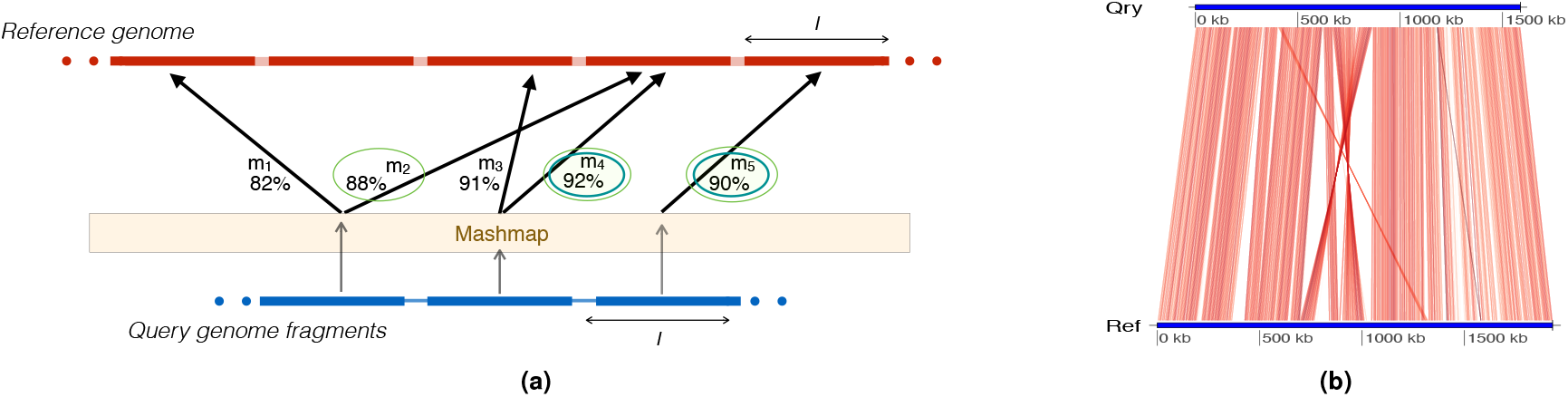
**a.** Graphical illustration of FastANI’s work-flow for computing ANI between a query genome and a reference genome. Five mappings are obtained from three query fragments using Mashmap (15). ***M**_forward_* saves the maximum identity mapping for each query fragment. In this example, ***M**_forward_* = {*m*_2_, *m*_4_, *m*_5_}. this set, *M_reciproca_ı* picks *m*_4_ and *m*_5_ as the maximum identity mapping for each reference bin. Mapping identities of orthologous mappings, thus found in ***M**_reciprocal_*, are finally averaged to compute ANI. **b.** FastANI supports visualization of the orthologous mappings ***M**_reciprocal_* that are used to estimate the ANI value using genoPlotR (25). In this figure, ANI is computed between *Bartonella quintana* strain (NC_018533.1) as query and *Bartonella henselae* strain (NC_005956.1) as reference. Red line segments denote the orthologous mappings computed by FastANI for ANI estimation.

### The Mashmap Sequence Mapping Algorithm

Given a query sequence, Mashmap (15) finds all its mapping positions in the reference sequence(s) above a user specified minimum alignment identity cut-off ***I***_0_ with high probability. Mashmap avoids direct alignments, but instead relates alignment identity (***I***) between sequences ***A*** and ***B*** to Jaccard similarity of constituent k-mers (***J***) under the Poisson distribution model:

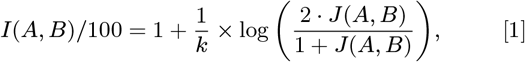

where *k* is the ***k***-mer size (16).

To estimate the Jaccard similarity itself, Mashmap uses a winnowed-MinHash estimator (15). This estimator requires only a small sample of k-mers from the query and reference sequences to be examined [see (15) for further details of Mashmap].

### FastANI Extends Mashmap to Compute ANI

Previously established and widely used implementations of ANI begin by either identifying the protein coding genomic fragments (8) or extracting approximately 1 Kbp long overlapping fragments from the query genome (3). These fragments are then mapped to the reference genome using BLASTn (7) or MUMmer (26), and the best match for each fragment is saved. This is followed by a reverse search, i.e., swapping the reference and query genomes. Mean identity of the reciprocal best matches computed through forward and reverse searches yields the ANI value. Rationale for this bi-directional approach is to bound the ANI computation to orthologous genes and discard the paralogs. In designing FastANI, we followed a similar approach while avoiding the alignment step.

FastANI first fragments the given query genome (***A***) into non-overlapping fragments of size ***l***. These ***l***-sized fragments are then mapped to the reference genome (***B***) using Mashmap. Mashmap first indexes the reference genome and subsequently computes mappings as well as alignment identity estimates for each query fragment, one at a time. At the end of the Mashmap run, all the query fragments 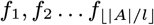 are mapped to ***B***. The results are saved in a set ***M*** containing triplets of the form 〈***f***, ***i***, ***p***⍪, where ***f*** is the fragment id, ***i*** is the identity estimate, and *p* is the starting position where ***f*** is mapped to **B**. The subset of ***M*** (say ***M_forward_***) corresponding to the maximum identity mapping for each query fragment is then extracted. To further identify the reciprocal matches, each triplet 〈***f***, ***i***, ***p***〉 in ***M_forward_*** is ‘binned’ based on its mapping position in the reference, with its value updated to 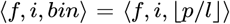. Through this step, fragments which are mapped to the same or nearby positions on the reference genome are likely to get equal bin value. Next, ***M_reciprocal_*** filters the maximum identity mapping for each bin. Finally, FastANI reports the mean identity of all the triplets in ***M_reciprocal_*** (See Figure 5 for an example and visualization).

We define ***τ*** as an input parameter to FastANI to indicate a minimum count of reciprocal mappings for the resulting ANI value to be trusted. It is important to appropriately choose the parameters (***l***, ***τ*** and ***I***_0_).

### FastANI Algorithm Parameter Settings

FastANI is targeted to estimate ANI in the 80%-100% identity range. Therefore, it calls Mashmap mapping routine with an identity cutoff ***I***_0_ = 80%, which enables it to compute mappings with alignment identity close to 80% or higher.

Choosing an appropriate value of query fragment ***l*** requires an evaluation of the trade-off between FastANI’s computation efficiency and ANI’s estimation accuracy. Higher value of ***l*** implies less number of non-overlapping query fragments, thus reducing the overall runtime. However, if ***l*** is much longer than the average gene length, a fragment could span more than one conserved segment, especially if the genome is highly recombinant. We empirically evaluated different values of ***l*** and set it to 3 Kbp (Supplementary Table S1). Last, we set ***τ*** to 50 to avoid incorrect ANI estimation from just a few matching fragments between genomes that are too divergent (e.g., showing <80% ANI). With ***l*** = 3 Kbp, ***τ*** =50 implies that we require at least 150 Kbp homologous genome sequence between two genomes to make a reliable ANI estimate, which is a reasonable assumption for both complete and incomplete genome assemblies based on our previous study (27).

### Software and Data Availability

FastANI can be downloaded at https://github.com/ParBUSS/FastANI. All the datasets used in this study are available at http://enve-omics.ce.gatech.edu/data/fastani.

## ACKNOWLEDGMENTS

We thank Jim Cole for discussions regarding the k-mer component of the FastANI algorithm. This work was supported, in part, by US National Science Foundation awards 1356288 (to KTK) and IIS-1416259 (to SA).

## Supplementary material

**Table S1:** Evaluation of FastANI accuracy and performance while varying the fragment length *I* used in the algorithm. We measure Pearson correlation coefficients of FastANI estimate with BLAST-based ANI computation (ANI_b_) as well as runtime and memory usage for each value of fragment size (1 Kbp – 10 Kbp). This experiment was conducted using datasets D3 and D4. From the table, it is evident that increasing fragment size improves runtime and memory usage, but negatively affects accuracy. Based on these tradeoffs, we set the fragment size to 3 Kbp in the FastANI implementation.

| Dataset | Metric | Fragment size |  |  |  |
| --- | --- | --- | --- | --- | --- |
|  |  | 1 Kbp | 3 Kbp | 5 Kbp | 10 Kbp |
| D3 | Correlation with $ANI_b$ | 0.9980964 | 0.9952683 | 0.9919395 | 0.9867375 |
|  | Runtime (index phase) in seconds | 2589.72 | 1666.58 | 1435.32 | 1145.05 |
|  | Runtime (compute phase) in seconds | 4737.62 | 2099.08 | 1286.37 | 546.78 |
|  | Memory (GB) | 113.75 | 48.35 | 28.95 | 14.73 |
| D4<br>(without two outlier genome assemblies) | Correlation with $ANI_b$ | 0.9649292 | 0.9439152 | 0.9019418 | 0.7865688 |
|  | Runtime (index phase) in seconds | 256.63 | 175.20 | 158.73 | 124.71 |
|  | Runtime (compute phase) in seconds | 746.63 | 254.11 | 172.82 | 74.32 |
|  | Memory (GB) | 12.59 | 4.38 | 3.16 | 1.59 |

**Figure S1.**
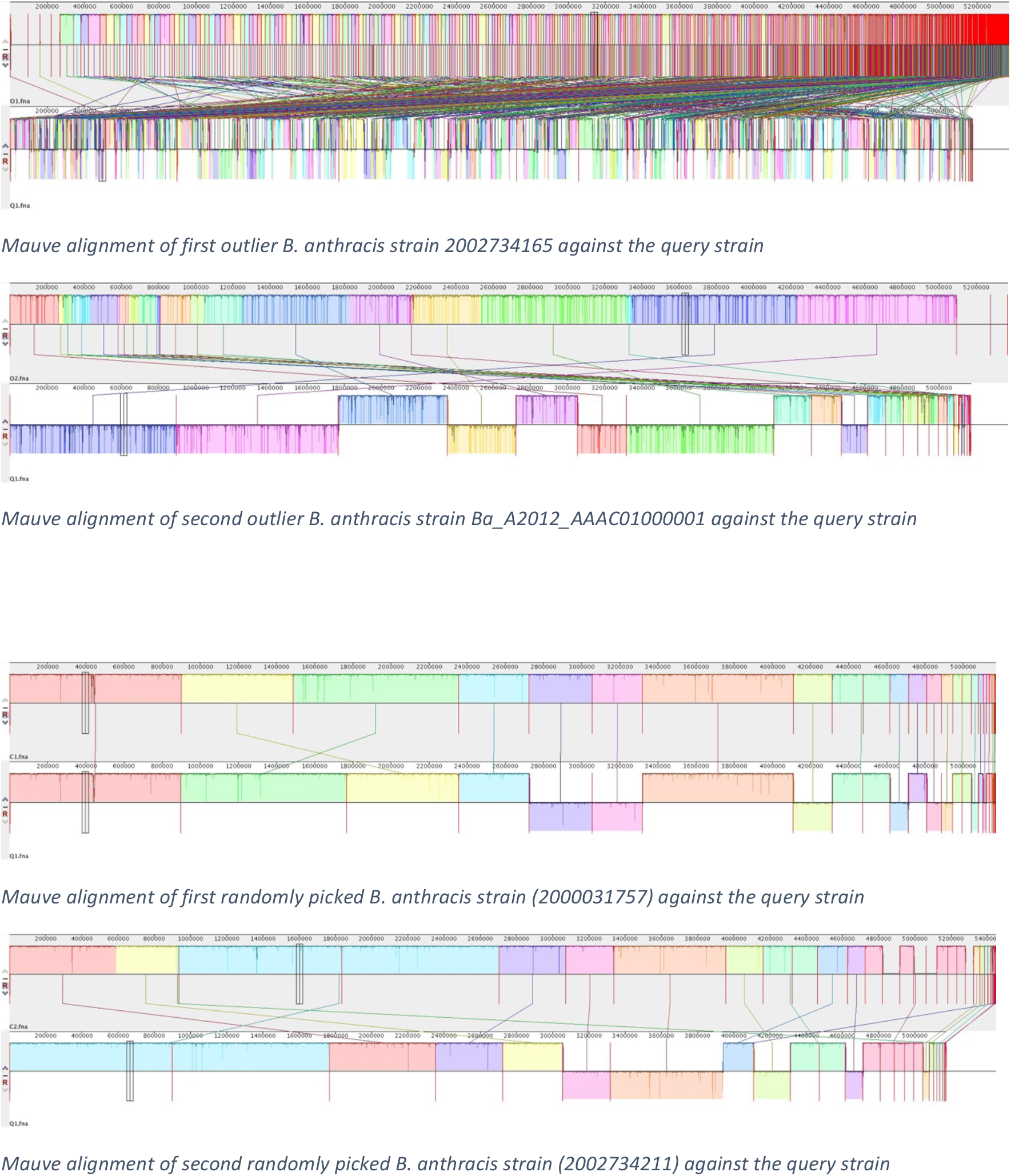
Top two plots show the mauve alignments of the two outlier *B. anthracis* strains (2002734165 and Ba_A2012_AAAC01000001) against the query strain (2000031001) used in D4 dataset. Bottom two plots show the mauve alignments of two randomly picked *B. anthracis* strains against the query strain. The top two outlier strains show unusually high degree of recombination and gaps than we expect between any two correctly sequenced and assembled *B. anthracis* strains. Same behavior was also observed using visualization support in FastANI software (figures not shown here).

**Figure S2.**
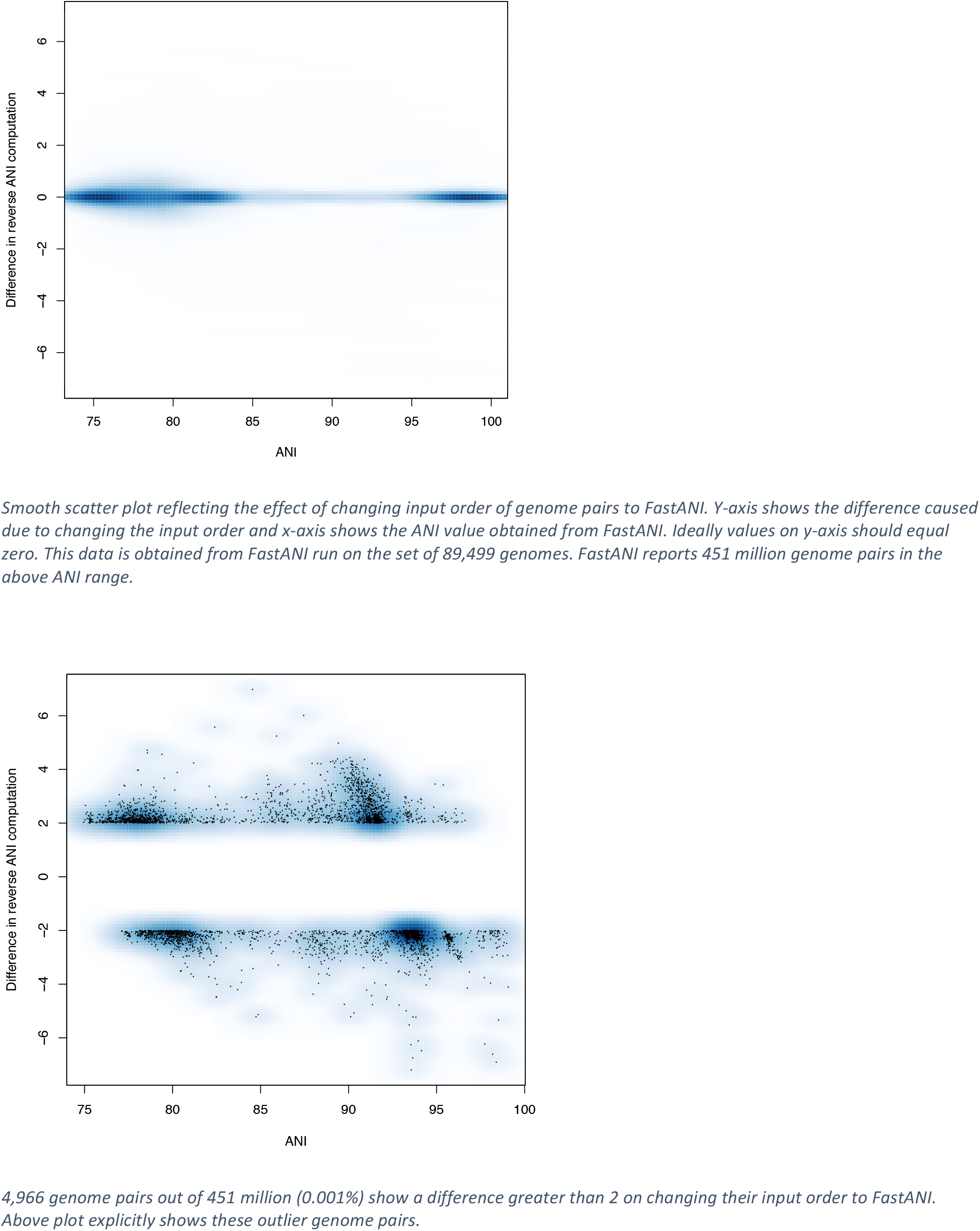
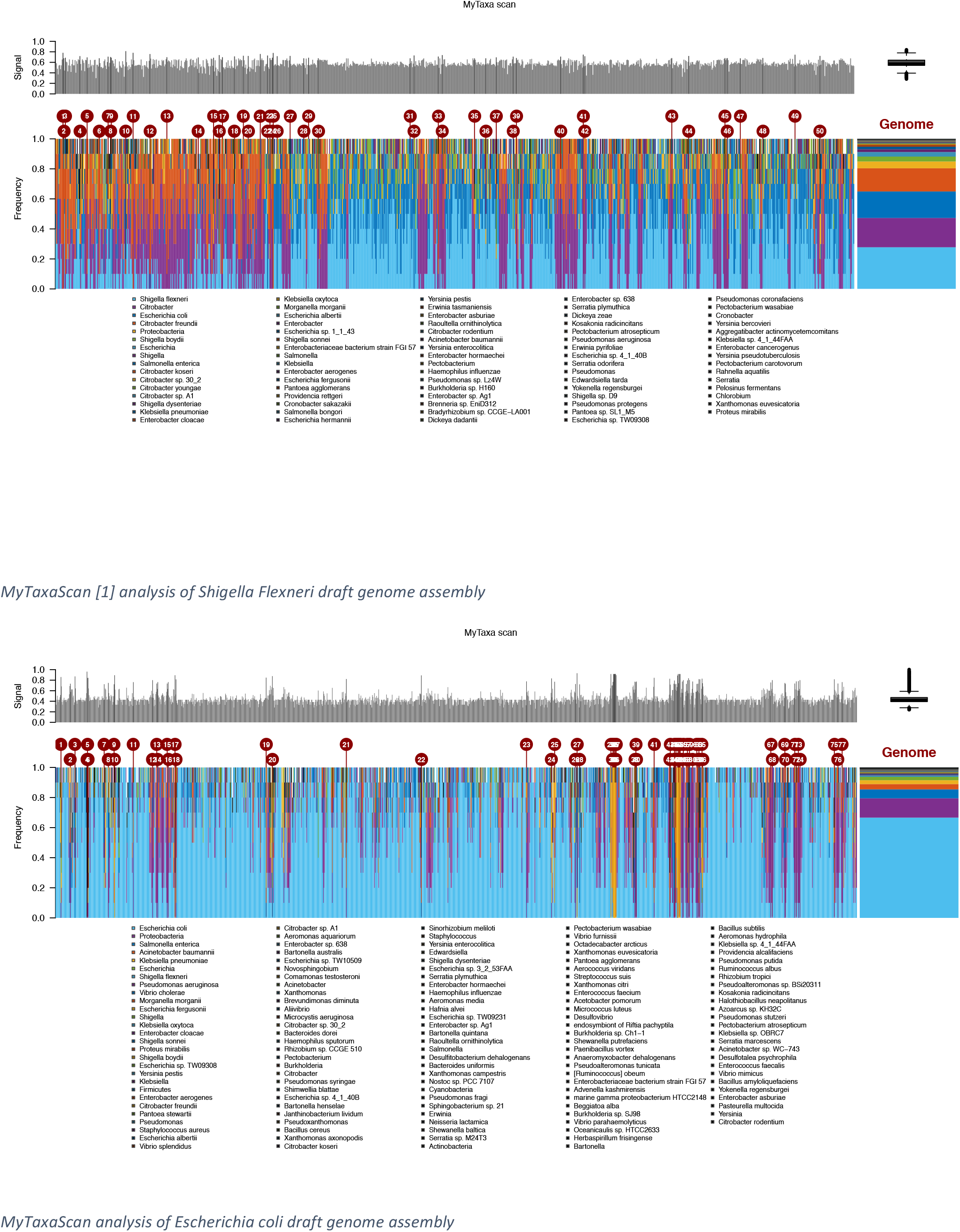
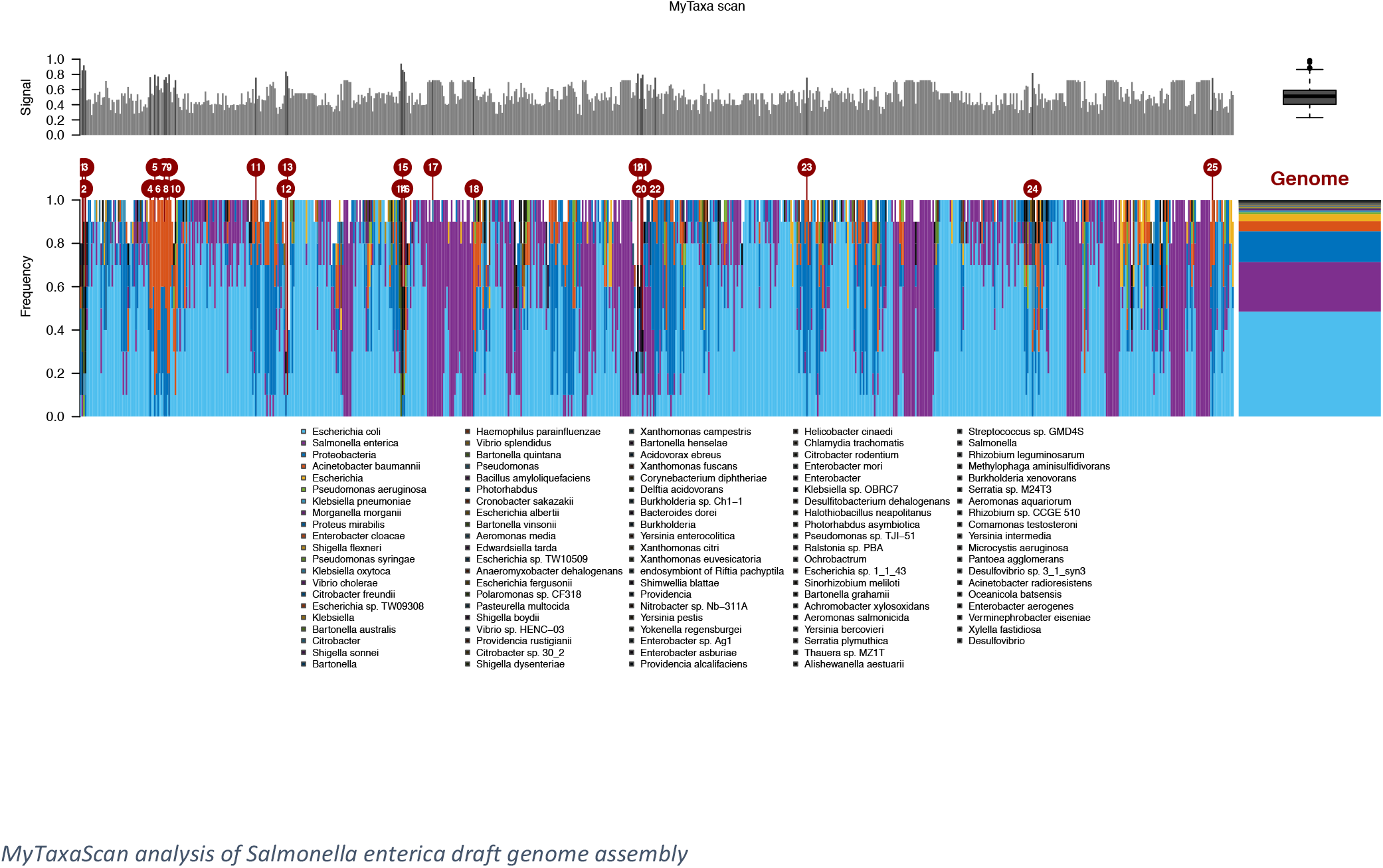
FastANI accepts genome pairs <reference genome, query genome> as input. For most genome pairs, input order causes an insignificant change in the FastANI tool’s ANI estimate. We first demonstrate this by showing a smooth-scatter plot of 451M genome pairs. Among them, we observe 4,966 outlier genome pairs (0.001% of 451M) that show difference ≥ 2. On further investigating these outliers, we conclude that they are caused by contaminated genome assemblies. We analyze top three genomes that contribute to the 4,966 outliers: a) Shigella_flexneri draft assembly (part of 1,941 outliers), b) Escherichia coli draft assembly (part of 632 outliers), and c) Salmonella enterica draft assembly (part of 505 outliers). Multiple colored peaks in all the three MyTaxaScan [1] plots highlight significant contamination in these assemblies from other species.

**Figure S3.**
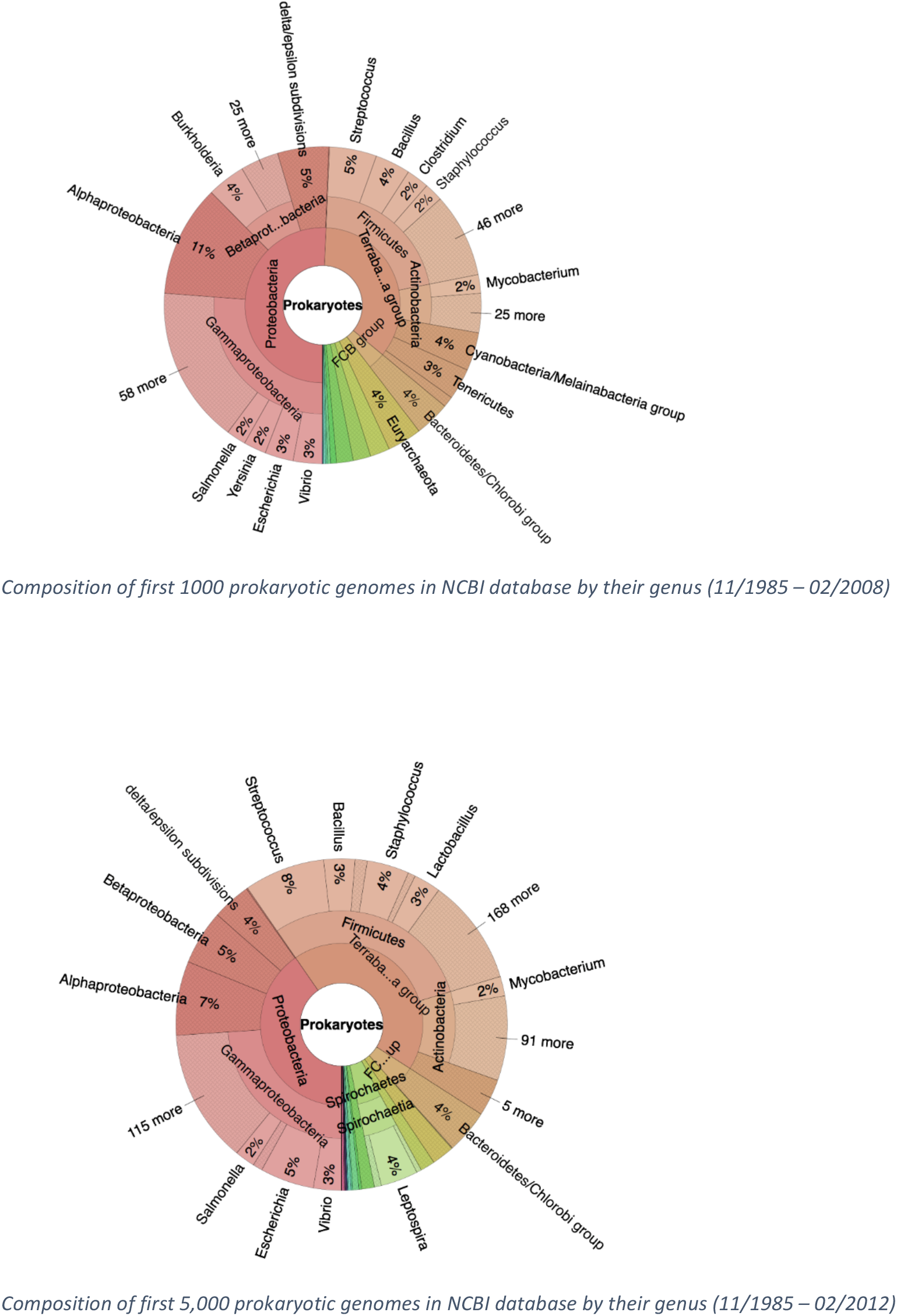
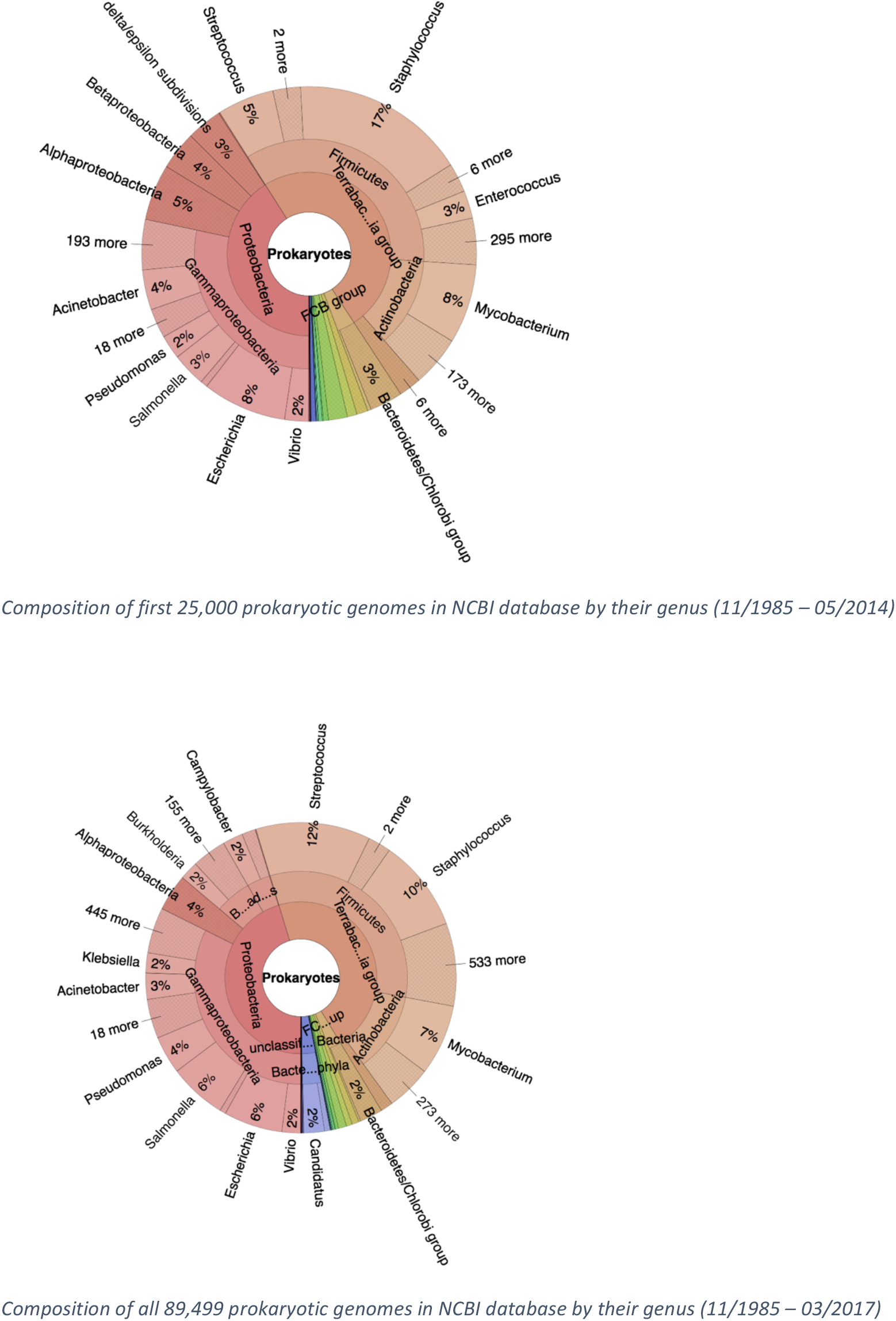
Composition of draft prokaryotic assemblies in the NCBI database with time at the genus level is visualized using Krona [2] charts. As expected, genus of high known biological significance started to dominate the database progressively. For each of these cohorts, ANI distribution density curves are shown in Fig. 5b.

